## Supplementary material for "Whole genome association testing in 333,100 individuals across three biobanks identifies rare non-coding single variant and genomic aggregate associations with height": TOPMed Acknowledgements

### INVESTIGATOR ACKNOWLEDGEMENTS

- Xihao Li was funded in part by NHLBI TOPMed Fellowship.
- Sharon L.R. Kardia was funded in part by NIH R01 HL119443, 3R01HL055673-18S1, HHSN268201500014C.
- Nicholette D. Palmer was funded in part by NIH R01AG058921.
- Donna K. Arnett was funded in part by NIH U01 HL072524 and R01 HL104135-04S1, U01 HL054472, U01 HL054473, U01 HL054495, and U01 HL054509, R01 HL055673 with supplement-18S1.
- Allison E. Ashley-Koch was funded in part by NIH R01HL68959.
- Jiang He was funded in part by NIH U01HL072507, P20GM109036.
- Ruth J.F. Loos was funded in part by NIH R01HL142302; R01DK124097; R01 DK110113; R01DK107786; X01HL134588.
- Bruce M. Psaty was funded in part by HL105756.
- Edwin K. Silverman was funded in part by U01HL089856, R01HL152728, R01HL133135, R01HL147148, and P01HL114501.
- Scott T. Weiss was funded in part by P01 HL132825.
- Ching-Ti Liu was funded in part by NIH NIDDK R01DK122503.
- Kari E. North was funded in part by R01HG010297; U01HG007416, DK122503.
- Anne E. Justice was funded in part by NIH NIDDK R01 DK 122503.
- Xihong Lin was funded in part by R35-CA197449, P01-CA134294, U19-CA203654, R01-HL113338, and U01-HG009088.

### STUDY ACKNOWLEDGEMENTS

**NHLBI TOPMed Informatics Research Center and Data Coordinating Center:**

Core support including centralized genomic read mapping and genotype calling, along with variant quality metrics and filtering were provided by the TOPMed Informatics Research Center (3R01HL-117626-02S1; contract HHSN268201800002I). Core support including phenotype harmonization, data management, sample-identity QC, and general program coordination were provided by the TOPMed Data Coordinating Center (R01HL-120393; U01HL-120393; contract HHSN268201800001I). We gratefully acknowledge the studies and participants who provided biological samples and data for TOPMed.

**NHLBI TOPMed:** **Genetics of Cardiometabolic Health in the Amish (Amish)**

Genome sequencing for “NHLBI TOPMed: Whole Genome Sequencing and Related Phenotypes in the Genetics of Cardiometabolic Health in the Amish” (phs000956) was performed at the Broad Institute Genomics Platform (3R01HL121007-01S1).

**NHLBI TOPMed:** **Atherosclerosis Risk in Communities Study VTE cohort (ARIC)**

The Atherosclerosis Risk in Communities study has been funded in whole or in part with Federal funds from the National Heart, Lung, and Blood Institute, National Institutes of Health, Department of Health and Human Services (contract numbers HHSN268201700001I, HHSN268201700002I, HHSN268201700003I, HHSN268201700004I and HHSN268201700005I). The authors thank the staff and participants of the ARIC study for their important contributions.

Genome sequencing for “NHLBI TOPMed: Whole Genome Sequencing and Related Phenotypes in the Atherosclerosis Risk in Communities Study VTE cohort” (phs001211) was performed at the Baylor College of Medicine Human Genome Sequencing Center (3U54HG003273-12S2, HHSN268201500015C).

**NHLBI TOPMed:** **New Approaches for Empowering Studies of Asthma in Populations of African Descent - Barbados Asthma Genetics Study (BAGS)**

We gratefully acknowledge the contributions of Pissamai and Trevor Maul, Paul Levett, Anselm Hennis, P. Michele Lashley, Raana Naidu, Malcolm Howitt and Timothy Roach, and the numerous health care providers, and community clinics and co-investigators who assisted in the phenotyping and collection of DNA samples, and the families and patients for generously donating DNA samples to the Barbados Asthma Genetics Study (BAGS). Funding for BAGS was provided by National Institutes of Health (NIH) R01HL104608, R01HL087699, and HL104608 S1.

Genome sequencing for “NHLBI TOPMed: New Approaches for Empowering Studies of Asthma in Populations of African Descent - Barbados Asthma Genetics Study” (phs001143) was performed at Illumina (3R01HL104608-04S1).

**NHLBI TOPMed:** **Mount Sinai BioMe Biobank (BioMe)**

The Mount Sinai BioMe Biobank has been supported by The Andrea and Charles Bronfman Philanthropies and in part by Federal funds from the NHLBI and NHGRI (U01HG00638001; U01HG007417; X01HL134588). We thank all participants in the Mount Sinai Biobank. We also thank all our recruiters who have assisted and continue to assist in data collection and management and are grateful for the computational resources and staff expertise provided by Scientific Computing at the Icahn School of Medicine at Mount Sinai.

Genome sequencing for “NHLBI TOPMed: Mount Sinai BioMe Biobank” (phs001644) was performed at the Baylor College of Medicine Human Genome Sequencing Center (HHSN268201600033I) and McDonnell Genome Institute (HHSN268201600037I).

**NHLBI TOPMed:** **Coronary Artery Risk Development in Young Adults (CARDIA)**

The Coronary Artery Risk Development in Young Adults Study (CARDIA) is conducted and supported by the National Heart, Lung, and Blood Institute (NHLBI) in collaboration with the University of Alabama at Birmingham (HHSN268201800005I & HHSN268201800007I), Northwestern University (HHSN268201800003I), University of Minnesota (HHSN268201800006I), and Kaiser Foundation Research Institute (HHSN268201800004I). CARDIA was also partially supported by the Intramural Research Program of the National Institute on Aging (NIA) and an intra‐agency agreement between NIA and NHLBI (AG0005).

Genome sequencing for “NHLBI TOPMed: Coronary Artery Risk Development in Young Adults” (phs001612) was performed at the Baylor College of Medicine Human Genome Sequencing Center (HHSN268201600033I).

**NHLBI TOPMed:** **Cleveland Clinic Atrial Fibrillation Study (CCAF)**

This study was supported by the National Institutes of Health (NIH) grants R01 HL 090620 and R01 HL 111314, the NIH National Center for Research Resources for Case Western Reserve University and Cleveland Clinic Clinical and Translational Science Award UL1-RR024989, the Cleveland Clinic Department of Cardiovascular Medicine philanthropy research funds, and the Tomsich Atrial Fibrillation Research Fund.

Genome sequencing for “NHLBI TOPMed: Cleveland Clinic Atrial Fibrillation Study” (phs001189) was performed at the Broad Institute Genomics Platform (3R01HL092577-06S1).

**NHLBI TOPMed: Cleveland Family Study - WGS Collaboration (CFS)**

The Cleveland Family Study has been supported in part by National Institutes of Health grants [R01-HL046380, KL2-RR024990, R35-HL135818, and R01-HL113338].

Genome sequencing for “NHLBI TOPMed: Cleveland Family Study - WGS Collaboration” (phs000954) was performed at the Northwest Genomics Center (3R01HL098433-05S1, HHSN268201600032I).

**NHLBI TOPMed:** **Cardiovascular Health Study (CHS)**

Cardiovascular Health Study: This research was supported by contracts HHSN268201200036C, HHSN268200800007C, HHSN268201800001C, N01HC55222, N01HC85079, N01HC85080, N01HC85081, N01HC85082, N01HC85083, N01HC85086, 75N92021D00006, and grants R01HL105756, U01HL080295 and U01HL130114 from the National Heart, Lung, and Blood Institute (NHLBI), with additional contribution from the National Institute of Neurological Disorders and Stroke (NINDS). Additional support was provided by R01AG023629 from the National Institute on Aging (NIA). A full list of principal CHS investigators and institutions can be found at CHS-NHLBI.org. The content is solely the responsibility of the authors and does not necessarily represent the official views of the National Institutes of Health.

Genome sequencing for “NHLBI TOPMed: Cardiovascular Health Study” (phs000954) was performed at the Broad Institute Genomics Platform (HHSN268201600034I) and the Baylor College of Medicine Human Genome Sequencing Center (HHSN268201600033I).

**NHLBI TOPMed:** **Genetic Epidemiology of COPD Study (COPDGene)**

The COPDGene project described was supported by Award Number U01 HL089897 and Award Number U01 HL089856 from the National Heart, Lung, and Blood Institute. The content is solely the responsibility of the authors and does not necessarily represent the official views of the National Heart, Lung, and Blood Institute or the National Institutes of Health. The COPDGene project is also supported by the COPD Foundation through contributions made to an Industry Advisory Board that has included AstraZeneca, Bayer Pharmaceuticals, Boehringer Ingelheim, Genentech, GlaxoSmithKline, Novartis, Pfizer, Siemens and Sunovion. A full listing of COPDGene investigators can be found at: <http://www.copdgene.org/directory>.

Genome sequencing for “NHLBI TOPMed: Genetic Epidemiology of COPD Study” (phs000951) was performed at the Broad Institute Genomics Platform (HHSN268201500014C) and the Northwest Genomics Center (3R01HL089856-08S1).

**COPDGene® Investigators – Core Units:** Administrative Center: James D. Crapo, MD (PI); Edwin K. Silverman, MD, PhD (PI); Barry J. Make, MD; Elizabeth A. Regan, MD, PhD. ***Genetic Analysis Center:*** Terri Beaty, PhD; Ferdouse Begum, PhD; Peter J. Castaldi, MD, MSc; Michael Cho, MD; Dawn L. DeMeo, MD, MPH; Adel R. Boueiz, MD; Marilyn G. Foreman, MD, MS; Eitan Halper-Stromberg; Lystra P. Hayden, MD, MMSc; Craig P. Hersh, MD, MPH; Jacqueline Hetmanski, MS, MPH; Brian D. Hobbs, MD; John E. Hokanson, MPH, PhD; Nan Laird, PhD; Christoph Lange, PhD; Sharon M. Lutz, PhD; Merry-Lynn McDonald, PhD; Margaret M. Parker, PhD; Dmitry Prokopenko, Ph.D; Dandi Qiao, PhD; Elizabeth A. Regan, MD, PhD; Phuwanat Sakornsakolpat, MD; Edwin K. Silverman, MD, PhD; Emily S. Wan, MD; Sungho Won, PhD. ***Imaging Center:*** Juan Pablo Centeno; Jean-Paul Charbonnier, PhD; Harvey O. Coxson, PhD; Craig J. Galban, PhD; MeiLan K. Han, MD, MS; Eric A. Hoffman, Stephen Humphries, PhD; Francine L. Jacobson, MD, MPH; Philip F. Judy, PhD; Ella A. Kazerooni, MD; Alex Kluiber; David A. Lynch, MB; Pietro Nardelli, PhD; John D. Newell, Jr., MD; Aleena Notary; Andrea Oh, MD; Elizabeth A. Regan, MD, PhD; James C. Ross, PhD; Raul San Jose Estepar, PhD; Joyce Schroeder, MD; Jered Sieren; Berend C. Stoel, PhD; Juerg Tschirren, PhD; Edwin Van Beek, MD, PhD; Bram van Ginneken, PhD; Eva van Rikxoort, PhD; Gonzalo Vegas Sanchez-Ferrero, PhD; Lucas Veitel; George R. Washko, MD; Carla G. Wilson, MS; ***PFT QA Center, Salt Lake City, UT:*** Robert Jensen, PhD. **Data Coordinating Center and *Biostatistics, National Jewish Health, Denver, CO:*** Douglas Everett, PhD; Jim Crooks, PhD; Katherine Pratte, PhD; Matt Strand, PhD; Carla G. Wilson, MS. **Epidemiology Core, University of Colorado *Anschutz Medical Campus, Aurora, CO:*** John E. Hokanson, MPH, PhD; Gregory Kinney, MPH, PhD; Sharon M. Lutz, PhD; Kendra A. Young, PhD. ***Mortality Adjudication Core:*** Surya P. Bhatt, MD; Jessica Bon, MD; Alejandro A. Diaz, MD, MPH; MeiLan K. Han, MD, MS; Barry Make, MD; Susan Murray, ScD; Elizabeth Regan, MD; Xavier Soler, MD; Carla G. Wilson, MS. ***Biomarker Core:*** Russell P. Bowler, MD, PhD; Katerina Kechris, PhD; Farnoush Banaei-Kashani, Ph.D. **COPDGene® Investigators – Clinical Centers: *Ann Arbor VA:*** Jeffrey L. Curtis, MD; Perry G. Pernicano, MD. ***Baylor College of Medicine, Houston, TX:*** Nicola Hanania, MD, MS; Mustafa Atik, MD; Aladin Boriek, PhD; Kalpatha Guntupalli, MD; Elizabeth Guy, MD; Amit Parulekar, MD. ***Brigham and Women’s Hospital, Boston, MA:*** Dawn L. DeMeo, MD, MPH; Alejandro A. Diaz, MD, MPH; Lystra P. Hayden, MD; Brian D. Hobbs, MD; Craig Hersh, MD, MPH; Francine L. Jacobson, MD, MPH; George Washko, MD. ***Columbia University, New York, NY:*** R. Graham Barr, MD, DrPH; John Austin, MD; Belinda D’Souza, MD; Byron Thomashow, MD. ***Duke University Medical Center, Durham, NC:*** Neil MacIntyre, Jr., MD; H. Page McAdams, MD; Lacey Washington, MD. ***Grady Memorial Hospital, Atlanta, GA:*** Eric Flenaugh, MD; Silanth Terpenning, MD. ***HealthPartners Research Institute, Minneapolis, MN:*** Charlene McEvoy, MD, MPH; Joseph Tashjian, MD. ***Johns Hopkins University, Baltimore, MD:*** Robert Wise, MD; Robert Brown, MD; Nadia N. Hansel, MD, MPH; Karen Horton, MD; Allison Lambert, MD, MHS; Nirupama Putcha, MD, MHS. ***Lundquist Institute for Biomedical Innovationat Harbor UCLA Medical Center, Torrance, CA:*** Richard Casaburi, PhD, MD; Alessandra Adami, PhD; Matthew Budoff, MD; Hans Fischer, MD; Janos Porszasz, MD, PhD; Harry Rossiter, PhD; William Stringer, MD. ***Michael E. DeBakey VAMC, Houston, TX:*** Amir Sharafkhaneh, MD, PhD; Charlie Lan, DO. ***Minneapolis VA:*** Christine Wendt, MD; Brian Bell, MD; Ken M. Kunisaki, MD, MS. ***National Jewish Health, Denver, CO:*** Russell Bowler, MD, PhD; David A. Lynch, MB. ***Reliant Medical Group, Worcester, MA:*** Richard Rosiello, MD; David Pace, MD. ***Temple University, Philadelphia, PA:*** Gerard Criner, MD; David Ciccolella, MD; Francis Cordova, MD; Chandra Dass, MD; Gilbert D’Alonzo, DO; Parag Desai, MD; Michael Jacobs, PharmD; Steven Kelsen, MD, PhD; Victor Kim, MD; A. James Mamary, MD; Nathaniel Marchetti, DO; Aditi Satti, MD; Kartik Shenoy, MD; Robert M. Steiner, MD; Alex Swift, MD; Irene Swift, MD; Maria Elena Vega-Sanchez, MD. ***University of Alabama, Birmingham, AL:*** Mark Dransfield, MD; William Bailey, MD; Surya P. Bhatt, MD; Anand Iyer, MD; Hrudaya Nath, MD; J. Michael Wells, MD. ***University of California, San Diego, CA:*** Douglas Conrad, MD; Xavier Soler, MD, PhD; Andrew Yen, MD. ***University of Iowa, Iowa City, IA:*** Alejandro P. Comellas, MD; Karin F. Hoth, PhD; John Newell, Jr., MD; Brad Thompson, MD. ***University of Michigan, Ann Arbor, MI:*** MeiLan K. Han, MD MS; Ella Kazerooni, MD MS; Wassim Labaki, MD MS; Craig Galban, PhD; Dharshan Vummidi, MD. ***University of Minnesota, Minneapolis, MN:*** Joanne Billings, MD; Abbie Begnaud, MD; Tadashi Allen, MD. ***University of Pittsburgh, Pittsburgh, PA:*** Frank Sciurba, MD; Jessica Bon, MD; Divay Chandra, MD, MSc; Joel Weissfeld, MD, MPH . ***University of Texas Health, San Antonio, San Antonio, TX:*** Antonio Anzueto, MD; Sandra Adams, MD; Diego Maselli-Caceres, MD; Mario E. Ruiz, MD; Harjinder Singh.

**NHLBI TOPMed:** **The Genetic Epidemiology of Asthma in Costa Rica - Asthma in Costa Rica cohort (CRA)**

Genome sequencing for “NHLBI TOPMed: The Genetic Epidemiology of Asthma in Costa Rica - Asthma in Costa Rica cohort” (phs000988) was performed at the Northwest Genomics Center (3R37HL066289-13S1, HHSN268201600032I).

**NHLBI TOPMed:** **Diabetes Heart Study (DHS)**

This work was supported by R01 HL92301, R01 HL67348, R01 NS058700, R01 AR48797, R01 DK071891, R01 AG058921, the General Clinical Research Center of the Wake Forest University School of Medicine (M01 RR07122, F32 HL085989), the American Diabetes Association, and a pilot grant from the Claude Pepper Older Americans Independence Center of Wake Forest University Health Sciences (P60 AG10484).

Genome sequencing for “NHLBI TOPMed: Diabetes Heart Study” (phs001412) was performed at the Broad Institute Genomics Platform (HHSN268201500014C).

**NHLBI TOPMed: Framingham Heart Study (FHS)**

The Framingham Heart Study (FHS) acknowledges the support of contracts NO1-HC-25195, HHSN268201500001I and 75N92019D00031 from the National Heart, Lung and Blood Institute and grant supplement R01 HL092577-06S1 for this research. We also acknowledge the dedication of the FHS study participants without whom this research would not be possible. Dr. Vasan is supported in part by the Evans Medical Foundation and the Jay and Louis Coffman Endowment from the Department of Medicine, Boston University School of Medicine.

Genome sequencing for “NHLBI TOPMed: Whole Genome Sequencing and Related Phenotypes in the Framingham Heart Study” (phs000974.v1.p1) was performed at the Broad Institute Genomics Platform (3R01HL092577-06S1, 3U54HG003067-12S2).

**The Johns Hopkins Genetic Study of Atherosclerosis Risk (GeneSTAR)** was supported by grants from the National Institutes of Health through the National Heart, Lung, and Blood Institute (HL49762, HL071025, U01HL72518, HL087698, HL092165, HL099747, K23HL105897, HL112064) and the National Institute of Nursing Research (NR0224103), by a grant from the National Center for Research Resources (M01-RR000052) to the Johns Hopkins General Clinical Research Center, and by a grant from the National Center for Research Resources and the National Center for Advancing Translational Sciences (UL1 RR 025005) to the Johns Hopkins Institute for Clinical and Translational Research.

Genome sequencing for “The Johns Hopkins Genetic Study of Atherosclerosis Risk” (phs001218) was performed at Psomagen (3R01HL112064-04S1) and Illumina (R01HL112064), and the Broad Institute Genomics Platform (HHSN268201500014C).

**NHLBI TOPMed: Genetic Epidemiology Network of Arteriopathy (GENOA)**

Support for GENOA was provided by the National Heart, Lung and Blood Institute (U01 HL054457, U01 HL054464, U01 HL054481, R01 HL119443, and R01 HL087660) of the National Institutes of Health. WGS for “NHLBI TOPMed: Genetic Epidemiology Network of Arteriopathy” (phs001345) was performed at the Mayo Clinic Genotyping Core, the DNA Sequencing and Gene Analysis Center at the University of Washington (3R01HL055673-18S1), and the Broad Institute (HHSN268201500014C) for their genotyping and sequencing services. We would like to thank the GENOA participants.

**NHLBI TOPMed: Genetic Epidemiology Network of Salt Sensitivity (GenSalt)**

The Genetic Epidemiology Network of Salt-Sensitivity (GenSalt) was supported by research grants (U01HL072507, R01HL087263, and R01HL090682) from the National Heart, Lung and Blood Institute, National Institutes of Health, Bethesda, MD.

Genome sequencing for “NHLBI TOPMed: Genetic Epidemiology Network of Salt Sensitivity” (phs001217) was performed at the Baylor College of Medicine Human Genome Sequencing Center (HHSN268201500015C).

**NHLBI TOPMed: Hispanic Community Health Study - Study of Latinos (HCHS_SOL)**

The Hispanic Community Health Study/Study of Latinos is a collaborative study supported by contracts from the National Heart, Lung, and Blood Institute (NHLBI) to the University of North Carolina (HHSN268201300001I / N01-HC-65233), University of Miami (HHSN268201300004I / N01-HC-65234), Albert Einstein College of Medicine (HHSN268201300002I / N01-HC-65235), University of Illinois at Chicago – HHSN268201300003I / N01-HC-65236 Northwestern Univ), and San Diego State University (HHSN268201300005I / N01-HC-65237). The following Institutes/Centers/Offices have contributed to the HCHS/SOL through a transfer of funds to the NHLBI: National Institute on Minority Health and Health Disparities, National Institute on Deafness and Other Communication Disorders, National Institute of Dental and Craniofacial Research, National Institute of Diabetes and Digestive and Kidney Diseases, National Institute of Neurological Disorders and Stroke, NIH Institution-Office of Dietary Supplements. All HCHS/SOL participants provided informed consent, and the study was approved by the Institutional Review Board of local field centers, coordinating center and laboratories.

Genome sequencing for “NHLBI TOPMed: Hispanic Community Health Study - Study of Latinos” (phs001395) was performed at the Baylor College of Medicine Human Genome Sequencing Center (HHSN268201600033I).

**NHLBI TOPMed: Heart and Vascular Health Study (HVH)**

The Heart and Vascular Health Study was supported by grants HL068986, HL085251, HL095080, and HL073410 from the National Heart, Lung, and Blood Institute.

Genome sequencing for “NHLBI TOPMed: Heart and Vascular Health Study” (phs000993) was performed at the Broad Institute Genomics Platform (3R01HL092577-06S1) and the Baylor College of Medicine Human Genome Sequencing Center (3U54HG003273-12S2 / HHSN268201500015C).

**NHLBI TOPMed:** **Hypertension Genetic Epidemiology Network (HyperGEN)**

The HyperGEN Study is part of the National Heart, Lung, and Blood Institute (NHLBI) Family Blood Pressure Program; collection of the data represented here was supported by grants U01 HL054472 (MN Lab), U01 HL054473 (DCC), U01 HL054495 (AL FC), and U01 HL054509 (NC FC). The HyperGEN: Genetics of Left Ventricular Hypertrophy Study was supported by NHLBI grant R01 HL055673 with whole-genome sequencing made possible by supplement -18S1.

Genome sequencing for “NHLBI TOPMed: Hypertension Genetic Epidemiology Network” (phs001293) was performed at the Northwest Genomics Center (3R01HL055673-18S1).

**NHLBI TOPMed: Jackson Heart Study (JHS)**

The Jackson Heart Study (JHS) is supported and conducted in collaboration with Jackson State University (HHSN268201800013I), Tougaloo College (HHSN268201800014I), the Mississippi State Department of Health (HHSN268201800015I) and the University of Mississippi Medical Center (HHSN268201800010I, HHSN268201800011I and HHSN268201800012I) contracts from the National Heart, Lung, and Blood Institute (NHLBI) and the National Institute on Minority Health and Health Disparities (NIMHD). The authors also wish to thank the staffs and participants of the JHS. The views expressed in this manuscript are those of the authors and do not necessarily represent the views of the National Heart, Lung, and Blood Institute; the National Institutes of Health; or the U.S. Department of Health and Human Services.

Genome sequencing for “NHLBI TOPMed: The Jackson Heart Study” (phs000964.v1.p1) was performed at the Northwest Genomics Center (HHSN268201100037C).

**NHLBI TOPMed:** **Lung Tissue Research Consortium (LTRC)**

Genome sequencing for “NHLBI TOPMed: Lung Tissue Research Consortium” (phs000964.v1.p1) was performed at the Broad Institute Genomics Platform (HHSN268201600034I).

**NHLBI TOPMed:** **Mayo Clinic Venous Thromboembolism Study (Mayo_VTE)**

Funded, in part, by grants from the National Institutes of Health, National Heart, Lung and Blood Institute (HL66216 and HL83141). the National Human Genome Research Institute (HG04735, HG06379), and research support provided by Mayo Foundation.

Genome sequencing for “NHLBI TOPMed: Mayo Clinic Venous Thromboembolism Study” (phs001402) was performed at the Baylor College of Medicine Human Genome Sequencing Center (3U54HG003273-12S2 / HHSN268201500015C).

**NHLBI TOPMed: Multi-Ethnic Study of Atherosclerosis (MESA)**

The Multi-Ethnic Study of Atherosclerosis is a National Heart, Lung and Blood Institute-sponsored, population-based investigation of subclinical cardiovascular disease and its progression^1^. A total of 6,814 individuals, aged 45 to 84 years, were recruited from six US communities (Baltimore City and County, MD; Chicago, IL; Forsyth County, NC; Los Angeles County, CA; New York, NY; and St. Paul, MN) between July 2000 and August 2002. Participants were excluded if they had physician-diagnosed cardiovascular disease prior to enrollment, including angina, myocardial infarction, heart failure, stroke or TIA, resuscitated cardiac arrest or a cardiovascular intervention (e.g., CABG, angioplasty, valve replacement, or pacemaker/defibrillator placement). Pre-specified recruitment plans identified four racial/ethnic groups (White European-American, African-American, Hispanic-American, and Chinese-American) for enrollment, with targeted oversampling of minority groups to enhance statistical power.

Whole genome sequencing (WGS) for the Trans-Omics in Precision Medicine (TOPMed) program was supported by the National Heart, Lung and Blood Institute (NHLBI). WGS for “NHLBI TOPMed: Multi-Ethnic Study of Atherosclerosis (MESA)” (phs001416.v3.p1) was performed at the Broad Institute of MIT and Harvard (3U54HG003067-13S1). Centralized read mapping and genotype calling, along with variant quality metrics and filtering were provided by the TOPMed Informatics Research Center (3R01HL-117626-02S1). Phenotype harmonization, data management, sample-identity QC, and general study coordination, were provided by the TOPMed Data Coordinating Center (3R01HL-120393-02S1), and TOPMed MESA Multi-Omics (HHSN2682015000031/HSN26800004). The MESA projects are conducted and supported by the National Heart, Lung, and Blood Institute (NHLBI) in collaboration with MESA investigators. Support for the Multi-Ethnic Study of Atherosclerosis (MESA) projects are conducted and supported by the National Heart, Lung, and Blood Institute (NHLBI) in collaboration with MESA investigators. Support for MESA is provided by contracts 75N92020D00001, HHSN268201500003I, N01-HC-95159, 75N92020D00005, N01-HC-95160, 75N92020D00002, N01-HC-95161, 75N92020D00003, N01-HC-95162, 75N92020D00006, N01-HC-95163, 75N92020D00004, N01-HC-95164, 75N92020D00007, N01-HC-95165, N01-HC-95166, N01-HC-95167, N01-HC-95168, N01-HC-95169, UL1-TR-000040, UL1-TR-001079, UL1-TR-001420, UL1TR001881, DK063491, and R01HL105756. The authors thank the other investigators, the staff, and the participants of the MESA study for their valuable contributions. A full list of participating MESA investigators and institutes can be found at <http://www.mesa-nhlbi.org>.

**NHLBI TOPMed:** **Massachusetts General Hospital Atrial Fibrillation Study (MGH_AF)**

Genome sequencing for “NHLBI TOPMed: Massachusetts General Hospital Atrial Fibrillation Study” (phs001062) was performed at the Broad Institute Genomics Platform (3U54HG003067-12S2 / 3U54HG003067-13S1; 3U54HG003067-12S2 / 3U54HG003067-13S1; 3UM1HG008895-01S2, 3R01HL092577-06S1).

**NHLBI TOPMed:** **Outcome Modifying Genes in Sickle Cell Disease (OMG_SCD)**

The OMG-SCD study was administrated by Marilyn J. Telen, M.D. and Allison E. Ashley-Koch, Ph.D. from Duke University Medical Center and collection of the data set was supported by grants HL068959 and HL079915 from the National Heart, Lung, and Blood Institute (NHLBI) of the National Institute of Health (NIH).

Genome sequencing for “NHLBI TOPMed: Outcome Modifying Genes in Sickle Cell Disease” (phs001608) was performed at the Baylor College of Medicine Human Genome Sequencing Center (HHSN268201600033I, HHSN268201500015C).

**NHLBI TOPMed: Partners Healthcare Biorepository (Partners)**

Genome sequencing for “NHLBI TOPMed: Partners Healthcare Biorepository” (phs001024) was performed at the Broad Institute Genomics Platform (3R01HL092577-06S1).

**NHLBI TOPMed:** **Whole Genome Sequencing to Identify Causal Genetic Variants Influencing CVD Risk - San Antonio Family Studies (SAFS)**

Collection of the San Antonio Family Study data was supported in part by National Institutes of Health (NIH) grants R01 HL045522, MH078143, MH078111 and MH083824; and whole genome sequencing of SAFS subjects was supported by U01 DK085524 and R01 HL113323. We are very grateful to the participants of the San Antonio Family Study for their continued involvement in our research programs.

Genome sequencing for “NHLBI TOPMed: Whole Genome Sequencing to Identify Causal Genetic Variants Influencing CVD Risk - San Antonio Family Studies” (phs001215) was performed at Illumina (R01HL113322, 3R01HL113323-03S1).

**NHLBI TOPMed:** **Samoan Adiposity Study (Samoan)**

Date collection was funded by NIH grant R01-HL093093. We thank the Samoan participants of the study and local village authorities. We acknowledge the support of the Samoan Ministry of Health and the Samoa Bureau of Statistics for their support of this research.

Genome sequencing for “NHLBI TOPMed: Samoan Adiposity Study” (phs000972) was performed at the New York Genome Center Genomics (HHSN268201500016C) and the Northwest Genomics Center (HHSN268201100037C).

We would also like to acknowledge the Samoan Obesity, Lifestyle and Genetic Adaptations Study

(OLaGA) Group: Ranjan Deka, Dept. of Environmental Health, University of Cincinnati; Nicola L.

Hawley, Dept. of Chronic Disease Epidemiology, Yale University; Stephen T McGarvey, Dept. of

Epidemiology and International Health Institute, and Dept. of Anthropology, Brown University;

Ryan L Minster, Dept. of Human Genetics, University of Pittsburgh; Take Naseri, Ministry of

Health, Government of Samoa; Muagututi‘a Sefuiva Reupena, Lutia I Puava Ae Mapu I Fagalele;

Daniel E. Weeks, Depts. of Human Genetics and Biostatistics, University of Pittsburgh.

**NHLBI TOPMed: Taiwan Study of Hypertension using Rare Variants (THRV)**

The Rare Variants for Hypertension in Taiwan Chinese (THRV) is supported by the National Heart, Lung, and Blood Institute (NHLBI) grant (R01HL111249) and its participation in TOPMed is supported by an NHLBI supplement (R01HL111249-04S1). THRV is a collaborative study between Washington University in St. Louis, LA BioMed at Harbor UCLA, University of Texas in Houston, Taichung Veterans General Hospital, Taipei Veterans General Hospital, Tri-Service General Hospital, National Health Research Institutes, National Taiwan University, and Baylor University. THRV is based (substantially) on the parent SAPPHIRe study, along with additional population-based and hospital-based cohorts. SAPPHIRe was supported by NHLBI grants (U01HL54527, U01HL54498) and Taiwan funds, and the other cohorts were supported by Taiwan funds.

Genome sequencing for “NHLBI TOPMed: Taiwan Study of Hypertension using Rare Variants” (phs001387) was performed at the Baylor College of Medicine Human Genome Sequencing Center (3R01HL111249-04S1 / HHSN26820150015C).

**NHLBI TOPMed: Vanderbilt Atrial Fibrillation Ablation Registry (VAFAR)**

Genome sequencing for “NHLBI TOPMed: Vanderbilt Atrial Fibrillation Ablation Registry” (phs000997) was performed at the Broad Institute Genomics Platform (3R01HL092577-06S1, 3U54HG003067-12S2 / 3U54HG003067-13S1; 3UM1HG008895-01S2; 3UM1HG008895-01S2).

**NHLBI TOPMed:** **Vanderbilt Genetic Basis of Atrial Fibrillation (VU_AF)**

Genome sequencing for “NHLBI TOPMed: Vanderbilt Genetic Basis of Atrial Fibrillation” (phs001032) was performed at the Broad Institute Genomics Platform (3R01HL092577-06S1).

**NHLBI TOPMed:** **Treatment of Pulmonary Hypertension and Sickle Cell Disease with Sildenafil Therapy (walk_PHaSST)**

We thank Dr. Mark Gladwin and the investigators of the Walk-PHasst study and the patients who participated in the study. We also thanks the walk-PHaSST clinical site team: Albert Einstein College of Medicine: Jane Little and Verlene Davis; Columbia University: Robyn Barst, Erika Rosenzweig, Margaret Lee and Daniela Brady; UCSF Benioff Children's Hospital Oakland: Claudia Morris, Ward Hagar, Lisa Lavrisha, Howard Rosenfeld, and Elliott Vichinsky; Children’s Hospital of Pittsburgh of UPMC: Regina McCollum; Hammersmith Hospital, London: Sally Davies, Gaia Mahalingam, Sharon Meehan, Ofelia Lebanto, and Ines Cabrita; Howard University: Victor Gordeuk, Oswaldo Castro, Onyinye Onyekwere, Vandana Sachdev, Alvin Thomas, Gladys Onojobi, Sharmin Diaz, Margaret Fadojutimi-Akinsiku, and Randa Aladdin; Johns Hopkins University: Reda Girgis, Sophie Lanzkron and Durrant Barasa; NHLBI: Mark Gladwin, Greg Kato, James Taylor, Wynona Coles, Catherine Seamon, Mary Hall, Amy Chi, Cynthia Brenneman, Wen Li, and Erin Smith; University of Colorado: Kathryn Hassell, David Badesch, Deb McCollister and Julie McAfee; University of Illinois at Chicago: Dean Schraufnagel, Robert Molokie, George Kondos, Patricia Cole-Saffold, and Lani Krauz; National Heart & Lung Institute, Imperial College London: Simon Gibbs. Thanks also to the data coordination center team from Rho, Inc.: Nancy Yovetich, Rob Woolson, Jamie Spencer, Christopher Woods, Karen Kesler, Vickie Coble, and Ronald W. Helms. We also thank Dr. Yingze Zhang for directing the Walk-PHasst repository and Dr. Mehdi Nouraie for maintaining the Walk-PHasst database and Dr. Jonathan Goldsmith as a NIH program director for this study. Special thanks to the volunteers who participated in the Walk-PHaSST study. This project was funded with federal funds from the NHLBI, NIH, Department of Health and Human Services, under contract HHSN268200617182C. This study is registered at www.clinicaltrials.gov as NCT00492531. Detail description of the study was published in Blood, 2011 118:855-864, Machado et al "Hospitalization for pain in patients with sickle cell disease treated with sildenafil for elevated TRV and low exercise capacity".

Genome sequencing for “NHLBI TOPMed: Treatment of Pulmonary Hypertension and Sickle Cell Disease with Sildenafil Therapy” (phs001514) was performed at the Baylor College of Medicine Human Genome Sequencing Center (HHSN268201500015C).

**NHLBI TOPMed: Women's Genome Health Study (WGHS)**

The WGHS is supported by the National Heart, Lung, and Blood Institute (HL043851 and HL080467) and the National Cancer Institute (CA047988 and UM1CA182913). The most recent cardiovascular endpoints were supported by ARRA funding HL099355.

Genome sequencing for “NHLBI TOPMed: Women's Genome Health Study” (phs001040) was performed at the Broad Institute Genomics Platform (3R01HL092577-06S1).

**NHLBI TOPMed:** **Women's Health Initiative (WHI)**

The WHI program is funded by the National Heart, Lung, and Blood Institute, National Institutes of Health, U.S. Department of Health and Human Services through contracts 75N92021D00001, 75N92021D00002, 75N92021D00003, 75N92021D00004, 75N92021D00005.

Genome sequencing for “NHLBI TOPMed: Women's Health Initiative” (phs001237) was performed at the Broad Institute Genomics Platform (HHSN268201500014C).

**NHLBI TOPMed Consortium Banner Authors (in alphabetical order by last name)**

Abe, Namiko, New York Genome Center, New York, New York, 10013, US; Abecasis, Gonçalo,

University of Michigan, Ann Arbor, Michigan, 48109, US; Aguet, Francois, Broad Institute, Cambridge,

Massachusetts, 02142, US; Albert, Christine, Cedars Sinai, Boston, Massachusetts, 02114, US; Almasy,

Laura, Children's Hospital of Philadelphia, University of Pennsylvania, Philadelphia, Pennsylvania,

19104, US; Alonso, Alvaro, Emory University, Atlanta, Georgia, 30322, US; Ament, Seth, University of

Maryland, Baltimore, Maryland, 21201, US; Anderson, Peter, University of Washington, Seattle,

Washington, 98195, US; Anugu, Pramod, University of Mississippi, Jackson, Mississippi, 38677, US;

Applebaum-Bowden, Deborah, National Institutes of Health, Bethesda, Maryland, 20892, US; Ardlie,

Kristin, Broad Institute, Cambridge, Massachusetts, 02142, US; Arking, Dan, Johns Hopkins University,

Baltimore, Maryland, 21218, US; Arnett, Donna K, University of Kentucky, Lexington, Kentucky,

40506, US; Ashley-Koch, Allison, Duke University, Durham, North Carolina, 27708, US; Aslibekyan,

Stella, University of Alabama, Birmingham, Alabama, 35487, US; Assimes, Tim, Stanford University,

Stanford, California, 94305, US; Auer, Paul, Medical College of Wisconsin, Milwaukee, Wisconsin,

53211, US; Avramopoulos, Dimitrios, Johns Hopkins University, Baltimore, Maryland, 21218, US; Ayas,

Najib, Providence Health Care, Medicine, Vancouver, CA; Balasubramanian, Adithya, Baylor College of

Medicine Human Genome Sequencing Center, Houston, Texas, 77030, US; Barnard, John, Cleveland

Clinic, Cleveland, Ohio, 44195, US; Barnes, Kathleen, Tempus, University of Colorado Anschutz

Medical Campus, Aurora, Colorado, 80045, US; Barr, R. Graham, Columbia University, New York, New

York, 10032, US; Barron-Casella, Emily, Johns Hopkins University, Baltimore, Maryland, 21218, US;

Barwick, Lucas, The Emmes Corporation, LTRC, Rockville, Maryland, 20850, US; Beaty, Terri, Johns

Hopkins University, Baltimore, Maryland, 21218, US; Beck, Gerald, Cleveland Clinic, Quantitative

Health Sciences, Cleveland, Ohio, 44195, US; Becker, Diane, Johns Hopkins University, Medicine,

Baltimore, Maryland, 21218, US; Becker, Lewis, Johns Hopkins University, Baltimore, Maryland, 21218,

US; Beer, Rebecca, National Heart, Lung, and Blood Institute, National Institutes of Health, Bethesda,

Maryland, 20892, US; Beitelshees, Amber, University of Maryland, Baltimore, Maryland, 21201, US;

Benjamin, Emelia, Boston University, Massachusetts General Hospital, Boston University School of

Medicine, Boston, Massachusetts, 02114, US; Benos, Takis, University of Pittsburgh, Pittsburgh,

Pennsylvania, 15260, US; Bezerra, Marcos, Fundação de Hematologia e Hemoterapia de Pernambuco - Hemope, Recife, 52011-000, BR; Bielak, Larry, University of Michigan, Ann Arbor, Michigan, 48109,

US; Bis, Joshua, University of Washington, Cardiovascular Health Research Unit, Department of

Medicine, Seattle, Washington, 98195, US; Blackwell, Thomas, University of Michigan, Ann Arbor,

Michigan, 48109, US; Blangero, John, University of Texas Rio Grande Valley School of Medicine,

Human Genetics, Brownsville, Texas, 78520, US; Boerwinkle, Eric, University of Texas Health at

Houston, Houston, Texas, 77225, US; Bowden, Donald W., Wake Forest Baptist Health, Department of

Biochemistry, Winston-Salem, North Carolina, 27157, US; Bowler, Russell, National Jewish Health,

National Jewish Health, Denver, Colorado, 80206, US; Brody, Jennifer, University of Washington,

Seattle, Washington, 98195, US; Broeckel, Ulrich, Medical College of Wisconsin, Pediatrics, Milwaukee,

Wisconsin, 53226, US; Broome, Jai, University of Washington, Seattle, Washington, 98195, US; Brown,

Deborah, University of Texas Health at Houston, Pediatrics, Houston, Texas, 77030, US; Bunting, Karen,

New York Genome Center, New York, New York, 10013, US; Burchard, Esteban, University of

California, San Francisco, San Francisco, California, 94143, US; Bustamante, Carlos, Stanford

University, Biomedical Data Science, Stanford, California, 94305, US; Buth, Erin, University of

Washington, Biostatistics, Seattle, Washington, 98195, US; Cade, Brian, Brigham & Women's Hospital,

Brigham and Women's Hospital, Boston, Massachusetts, 02115, US; Cardwell, Jonathan, University of

Colorado at Denver, Denver, Colorado, 80204, US; Carey, Vincent, Brigham & Women's Hospital,

Boston, Massachusetts, 02115, US; Carrier, Julie, University of Montreal, , US; Carson, April, University

of Mississippi, Medicine, Jackson, Mississippi, 39213, US; Carty, Cara, Washington State University,

Pullman, Washington, 99164, US; Casaburi, Richard, University of California, Los Angeles, Los

Angeles, California, 90095, US; Casas Romero, Juan P, Brigham & Women's Hospital, , US; Casella,

James, Johns Hopkins University, Baltimore, Maryland, 21218, US; Castaldi, Peter, Brigham & Women's

Hospital, Medicine, Boston, Massachusetts, 02115, US; Chaffin, Mark, Broad Institute, Cambridge,

Massachusetts, 02142, US; Chang, Christy, University of Maryland, Baltimore, Maryland, 21201, US;

Chang, Yi-Cheng, National Taiwan University, Taipei, 10617, TW; Chasman, Daniel, Brigham &

Women's Hospital, Division of Preventive Medicine, Boston, Massachusetts, 02215, US; Chavan,

Sameer, University of Colorado at Denver, Denver, Colorado, 80204, US; Chen, Bo-Juen, New York

Genome Center, New York, New York, 10013, US; Chen, Wei-Min, University of Virginia,

Charlottesville, Virginia, 22903, US; Chen, Yii-Der Ida, Lundquist Institute, Torrance, California, 90502,

US; Cho, Michael, Brigham & Women's Hospital, Boston, Massachusetts, 02115, US; Choi, Seung Hoan,

Broad Institute, Cambridge, Massachusetts, 02142, US; Chuang, Lee-Ming, National Taiwan University,

National Taiwan University Hospital, Taipei, 10617, TW; Chung, Mina, Cleveland Clinic, Cleveland

Clinic, Cleveland, Ohio, 44195, US; Chung, Ren-Hua, National Health Research Institute Taiwan, Miaoli

County, 350, TW; Clish, Clary, Broad Institute, Metabolomics Platform, Cambridge, Massachusetts,

02142, US; Comhair, Suzy, Cleveland Clinic, Immunity and Immunology, Cleveland, Ohio, 44195, US;

Conomos, Matthew, University of Washington, Biostatistics, Seattle, Washington, 98195, US; Cornell,

Elaine, University of Vermont, Burlington, Vermont, 05405, US; Correa, Adolfo, University of

Mississippi, Population Health Science, Jackson, Mississippi, 39216, US; Crandall, Carolyn, University

of California, Los Angeles, Los Angeles, California, 90095, US; Crapo, James, National Jewish Health,

Denver, Colorado, 80206, US; Cupples, L. Adrienne, Boston University, Biostatistics, Boston,

Massachusetts, 02115, US; Curran, Joanne, University of Texas Rio Grande Valley School of Medicine,

Brownsville, Texas, 78520, US; Curtis, Jeffrey, University of Michigan, Internal Medicine, Ann Arbor,

Michigan, 48109, US; Custer, Brian, Vitalant Research Institute, San Francisco, California, 94118, US;

Damcott, Coleen, University of Maryland, Baltimore, Maryland, 21201, US; Darbar, Dawood, University

of Illinois at Chicago, Chicago, Illinois, 60607, US; David, Sean, University of Chicago, Chicago,

Illinois, 60637, US; Davis, Colleen, University of Washington, Seattle, Washington, 98195, US; Daya,

Michelle, University of Colorado at Denver, Denver, Colorado, 80204, US; de Andrade, Mariza, Mayo

Clinic, Health Quantitative Sciences Research , Rochester, Minnesota, 55905, US; de las Fuentes, Lisa,

Washington University in St Louis, Department of Medicine, Cardiovascular Division, St. Louis,

Missouri, 63110, US; de Vries, Paul, University of Texas Health at Houston, Human Genetics Center,

Department of Epidemiology, Human Genetics, and Environmental Sciences, Houston, Texas, 77030, US; DeBaun, Michael, Vanderbilt University, Nashville, Tennessee, 37235, US; Deka, Ranjan,

University of Cincinnati, Cincinnati, Ohio, 45220, US; DeMeo, Dawn, Brigham & Women's Hospital,

Boston, Massachusetts, 02115, US; Devine, Scott, University of Maryland, Baltimore, Maryland, 21201,

US; Dinh, Huyen, Baylor College of Medicine Human Genome Sequencing Center, Houston, Texas,

77030, US; Doddapaneni, Harsha , Baylor College of Medicine Human Genome Sequencing Center,

Houston, Texas, 77030, ; Duan, Qing, University of North Carolina, Chapel Hill, North Carolina, 27599,

US; Dugan-Perez, Shannon, Baylor College of Medicine Human Genome Sequencing Center, Houston,

Texas, 77030, US; Duggirala, Ravi, University of Texas Rio Grande Valley School of Medicine,

Edinburg, Texas, 78539, US; Durda, Jon Peter, University of Vermont, Burlington, Vermont, 05405, US;

Dutcher, Susan K., Washington University in St Louis, Genetics , St Louis, Missouri, 63110, US; Eaton,

Charles, Brown University, Providence, Rhode Island, 02912, US; Ekunwe, Lynette, University of

Mississippi, Jackson, Mississippi, 38677, US; El Boueiz, Adel, Harvard University, Channing Division of

Network Medicine, Cambridge, Massachusetts, 02138, US; Ellinor, Patrick, Massachusetts General

Hospital, Boston, Massachusetts, 02114, US; Emery, Leslie, University of Washington, Seattle,

Washington, 98195, US; Erzurum, Serpil, Cleveland Clinic, Cleveland, Ohio, 44195, US; Farber,

Charles, University of Virginia, Charlottesville, Virginia, 22903, US; Farek, Jesse, Baylor College of

Medicine Human Genome Sequencing Center, Houston, Texas, 77030, US; Fingerlin, Tasha, National

Jewish Health, Center for Genes, Environment and Health, Denver, Colorado, 80206, US; Flickinger,

Matthew, University of Michigan, Ann Arbor, Michigan, 48109, US; Fornage, Myriam, University of

Texas Health at Houston, Houston, Texas, 77225, US; Franceschini, Nora, University of North Carolina,

Epidemiology, Chapel Hill, North Carolina, 27599, US; Frazar, Chris, University of Washington, Seattle,

Washington, 98195, US; Fu, Mao, University of Maryland, Baltimore, Maryland, 21201, US; Fullerton,

Stephanie M., University of Washington, Seattle, Washington, 98195, US; Fulton, Lucinda, Washington

University in St Louis, St Louis, Missouri, 63130, US; Gabriel, Stacey, Broad Institute, Cambridge,

Massachusetts, 02142, US; Gan, Weiniu, National Heart, Lung, and Blood Institute, National Institutes of

Health, Bethesda, Maryland, 20892, US; Gao, Shanshan, University of Colorado at Denver, Denver,

Colorado, 80204, US; Gao, Yan, University of Mississippi, Jackson, Mississippi, 38677, US; Gass,

Margery, Fred Hutchinson Cancer Research Center, Seattle, Washington, 98109, US; Geiger, Heather,

New York Genome Center, New York City, New York, 10013, US; Gelb, Bruce, Icahn School of

Medicine at Mount Sinai, New York, New York, 10029, US; Geraci, Mark, University of Pittsburgh,

Pittsburgh, Pennsylvania, US; Germer, Soren, New York Genome Center, New York, New York, 10013,

US; Gerszten, Robert, Beth Israel Deaconess Medical Center, Boston, Massachusetts, 02215, US; Ghosh,

Auyon, Brigham & Women's Hospital, Boston, Massachusetts, 02115, US; Gibbs, Richard, Baylor

College of Medicine Human Genome Sequencing Center, Houston, Texas, 77030, US; Gignoux, Chris,

Stanford University, Stanford, California, 94305, US; Gladwin, Mark, University of Pittsburgh,

Pittsburgh, Pennsylvania, 15260, US; Glahn, David, Boston Children's Hospital, Harvard Medical School,

Department of Psychiatry, Boston, Massachusetts, 02115, US; Gogarten, Stephanie, University of

Washington, Seattle, Washington, 98195, US; Gong, Da-Wei, University of Maryland, Baltimore,

Maryland, 21201, US; Goring, Harald, University of Texas Rio Grande Valley School of Medicine, San

Antonio, Texas, 78229, US; Graw, Sharon, University of Colorado Anschutz Medical Campus, Aurora,

Colorado, 80045, US; Gray, Kathryn J., Mass General Brigham, Obstetrics and Gynecology, Boston,

Massachusetts, 02115, US; Grine, Daniel, University of Colorado at Denver, Denver, Colorado, 80204,

US; Gross, Colin, University of Michigan, Ann Arbor, Michigan, 48109, US; Gu, C. Charles, Washington

University in St Louis, St Louis, Missouri, 63130, US; Guan, Yue, University of Maryland, Baltimore,

Maryland, 21201, US; Guo, Xiuqing, Lundquist Institute, Torrance, California, 90502, US; Gupta,

Namrata, Broad Institute, Cambridge, Massachusetts, 02142, US; Haas, David M., Indiana University,

OB/GYN, Indianapolis, Indiana, 46202, US; Haessler, Jeff, Fred Hutchinson Cancer Research Center,

Seattle, Washington, 98109, US; Hall, Michael, University of Mississippi, Cardiology, Jackson,

Mississippi, 39216, US; Han, Yi, Baylor College of Medicine Human Genome Sequencing Center,

Houston, Texas, 77030, US; Hanly, Patrick, University of Calgary, Medicine, Calgary, CA; Harris,

Daniel, University of Maryland, Genetics, Philadelphia, Pennsylvania, 19104, US; Hawley, Nicola L., Yale University, Department of Chronic Disease Epidemiology, New Haven, Connecticut, 06520, US;

He, Jiang, Tulane University, New Orleans, Louisiana, 70118, US; Heavner, Ben, University of

Washington, Biostatistics, Seattle, Washington, 98195, US; Heckbert, Susan, University of Washington,

Epidemiology, Seattle, Washington, 98195, US; Hernandez, Ryan, University of California, San

Francisco, San Francisco, California, 94143, US; Herrington, David, Wake Forest Baptist Health,

Winston-Salem, North Carolina, 27157, US; Hersh, Craig, Brigham & Women's Hospital, Channing

Division of Network Medicine, Boston, Massachusetts, 02115, US; Hidalgo, Bertha, University of

Alabama, Birmingham, Alabama, 35487, US; Hixson, James, University of Texas Health at Houston,

Houston, Texas, 77225, US; Hobbs, Brian, Brigham & Women's Hospital, Boston, Massachusetts, 02115,

US; Hokanson, John, University of Colorado at Denver, Denver, Colorado, 80204, US; Hong, Elliott,

University of Maryland, Baltimore, Maryland, 21201, US; Hoth, Karin, University of Iowa, Iowa City,

Iowa, 52242, US; Hsiung, Chao (Agnes), National Health Research Institute Taiwan, Institute of

Population Health Sciences, NHRI, Miaoli County, 350, TW; Hu, Jianhong, Baylor College of Medicine

Human Genome Sequencing Center, Houston, Texas, 77030, US; Hung, Yi-Jen, Tri-Service General

Hospital National Defense Medical Center, , TW; Huston, Haley, Blood Works Northwest, Seattle,

Washington, 98104, US; Hwu, Chii Min, Taichung Veterans General Hospital Taiwan, Taichung City,

407, TW; Irvin, Marguerite Ryan, University of Alabama, Birmingham, Alabama, 35487, US; Jackson,

Rebecca, Oklahoma State University Medical Center, Internal Medicine, DIvision of Endocrinology,

Diabetes and Metabolism, Columbus, Ohio, 43210, US; Jain, Deepti, University of Washington, Seattle,

Washington, 98195, US; Jaquish, Cashell, National Heart, Lung, and Blood Institute, National Institutes

of Health, NHLBI, Bethesda, Maryland, 20892, US; Johnsen, Jill, Blood Works Northwest, Research

Institute, Seattle, Washington, 98104, US; Johnson, Andrew, National Heart, Lung, and Blood Institute,

National Institutes of Health, Bethesda, Maryland, 20892, US; Johnson, Craig, University of Washington,

Seattle, Washington, 98195, US; Johnston, Rich, Emory University, Atlanta, Georgia, 30322, US; Jones,

Kimberly, Johns Hopkins University, Baltimore, Maryland, 21218, US; Kang, Hyun Min, University of

Michigan, Biostatistics, Ann Arbor, Michigan, 48109, US; Kaplan, Robert, Albert Einstein College of

Medicine, New York, New York, 10461, US; Kardia, Sharon, University of Michigan, Ann Arbor,

Michigan, 48109, US; Kelly, Shannon, University of California, San Francisco, San Francisco,

California, 94118, US; Kenny, Eimear, Icahn School of Medicine at Mount Sinai, New York, New York,

10029, US; Kessler, Michael, University of Maryland, Baltimore, Maryland, 21201, US; Khan, Alyna,

University of Washington, Seattle, Washington, 98195, US; Khan, Ziad, Baylor College of Medicine

Human Genome Sequencing Center, Houston, Texas, 77030, US; Kim, Wonji, Harvard University,

Cambridge, Massachusetts, 02138, US; Kimoff, John, McGill University, Montréal, QC H3A 0G4, CA;

Kinney, Greg, University of Colorado at Denver, Epidemiology, Aurora, Colorado, 80045, US; Konkle,

Barbara, Blood Works Northwest, Medicine, Seattle, Washington, 98104, US; Kooperberg, Charles, Fred

Hutchinson Cancer Research Center, Seattle, Washington, 98109, US; Kramer, Holly, Loyola University,

Public Health Sciences, Maywood, Illinois, 60153, US; Lange, Christoph, Harvard School of Public

Health, Biostats, Boston, Massachusetts, 02115, US; Lange, Ethan, University of Colorado at Denver,

Denver, Colorado, 80204, US; Lange, Leslie, University of Colorado at Denver, Medicine, Aurora,

Colorado, 80048, US; Laurie, Cathy, University of Washington, Seattle, Washington, 98195, US; Laurie,

Cecelia, University of Washington, Seattle, Washington, 98195, US; LeBoff, Meryl, Brigham &

Women's Hospital, Boston, Massachusetts, 02115, US; Lee, Jiwon, Brigham & Women's Hospital,

Boston, Massachusetts, 02115, US; Lee, Sandra, Baylor College of Medicine Human Genome

Sequencing Center, Houston, Texas, 77030, US; Lee, Wen-Jane, Taichung Veterans General Hospital

Taiwan, Taichung City, 407, TW; LeFaive, Jonathon, University of Michigan, Ann Arbor, Michigan,

48109, US; Levine, David, University of Washington, Seattle, Washington, 98195, US; Levy, Dan,

National Heart, Lung, and Blood Institute, National Institutes of Health, Bethesda, Maryland, 20892, US;

Lewis, Joshua, University of Maryland, Baltimore, Maryland, 21201, US; Li, Xiaohui, Lundquist

Institute, Torrance, California, 90502, US; Li, Yun, University of North Carolina, Chapel Hill, North

Carolina, 27599, US; Lin, Henry, Lundquist Institute, Torrance, California, 90502, US; Lin, Honghuang,

Boston University, Boston, Massachusetts, 02215, US; Lin, Xihong, Harvard School of Public Health, Boston, Massachusetts, 02115, US; Liu, Simin, Brown University, Epidemiology and Medicine,

Providence, Rhode Island, 02912, US; Liu, Yongmei, Duke University, Cardiology, Durham, North

Carolina, 27708, US; Liu, Yu, Stanford University, Cardiovascular Institute, Stanford, California, 94305,

US; Loos, Ruth J.F., Icahn School of Medicine at Mount Sinai, The Charles Bronfman Institute for

Personalized Medicine, New York, New York, 10029, US; Lubitz, Steven, Massachusetts General

Hospital, Boston, Massachusetts, 02114, US; Lunetta, Kathryn, Boston University, Boston,

Massachusetts, 02215, US; Luo, James, National Heart, Lung, and Blood Institute, National Institutes of

Health, Bethesda, Maryland, 20892, US; Magalang, Ulysses, Ohio State University, Division of

Pulmonary, Critical Care and Sleep Medicine, Columbus, Ohio, 43210, US; Mahaney, Michael,

University of Texas Rio Grande Valley School of Medicine, Brownsville, Texas, 78520, US; Make,

Barry, Johns Hopkins University, Baltimore, Maryland, 21218, US; Manichaikul, Ani, University of

Virginia, Charlottesville, Virginia, 22903, US; Manning, Alisa, Broad Institute, Harvard University,

Massachusetts General Hospital, , ; Manson, JoAnn, Brigham & Women's Hospital, Boston,

Massachusetts, 02115, US; Martin, Lisa, George Washington University, cardiology, Washington,

District of Columbia, 20037, US; Marton, Melissa, New York Genome Center, New York City, New

York, 10013, US; Mathai, Susan, University of Colorado at Denver, Denver, Colorado, 80204, US;

Mathias, Rasika, Johns Hopkins University, Baltimore, Maryland, 21218, US; May, Susanne, University

of Washington, Biostatistics, Seattle, Washington, 98195, US; McArdle, Patrick, University of Maryland,

Baltimore, Maryland, 21201, US; McDonald, Merry-Lynn, University of Alabama, University of

Alabama at Birmingham, Birmingham, Alabama, 35487, US; McFarland, Sean, Harvard University,

Cambridge, Massachusetts, 02138, US; McGarvey, Stephen, Brown University, Epidemiology,

Providence, Rhode Island, 02912, US; McGoldrick, Daniel , University of Washington, Genome

Sciences, Seattle, Washington, 98195, US; McHugh, Caitlin, University of Washington, Biostatistics,

Seattle, Washington, 98195, US; McNeil, Becky, RTI International, , US; Mei, Hao, University of

Mississippi, Jackson, Mississippi, 38677, US; Meigs, James, Massachusetts General Hospital, Medicine,

Boston , Massachusetts, 02114, US; Menon, Vipin, Baylor College of Medicine Human Genome

Sequencing Center, Houston, Texas, 77030, US; Mestroni, Luisa, University of Colorado Anschutz

Medical Campus, Aurora, Colorado, 80045, US; Metcalf, Ginger, Baylor College of Medicine Human

Genome Sequencing Center, Houston, Texas, 77030, US; Meyers, Deborah A, University of Arizona,

Tucson, Arizona, 85721, US; Mignot, Emmanuel, Stanford University, Center For Sleep Sciences and

Medicine, Palo Alto, California, 94304, US; Mikulla, Julie, National Heart, Lung, and Blood Institute,

National Institutes of Health, Bethesda, Maryland, 20892, US; Min, Nancy, University of Mississippi,

Jackson, Mississippi, 38677, US; Minear, Mollie, National Institute of Child Health and Human

Development, National Institutes of Health, Bethesda, Maryland, 20892, US; Minster, Ryan L, University

of Pittsburgh, Pittsburgh, Pennsylvania, 15260, US; Mitchell, Braxton D., University of Maryland,

Baltimore, Maryland, 21201, US; Moll, Matt, Brigham & Women's Hospital, Medicine, Boston,

Massachusetts, 02115, US; Momin, Zeineen, Baylor College of Medicine Human Genome Sequencing

Center, Houston, Texas, 77030, US; Montasser, May E., University of Maryland, Baltimore, Maryland,

21201, US; Montgomery, Courtney, Oklahoma Medical Research Foundation, Genes and Human

Disease, Oklahoma City, Oklahoma, 73104, US; Muzny, Donna, Baylor College of Medicine Human

Genome Sequencing Center, Houston, Texas, 77030, US; Mychaleckyj, Josyf C, University of Virginia,

Charlottesville, Virginia, 22903, US; Nadkarni, Girish, Icahn School of Medicine at Mount Sinai, New

York, New York, 10029, US; Naik, Rakhi, Johns Hopkins University, Baltimore, Maryland, 21218, US;

Naseri, Take, Ministry of Health, Government of Samoa, Apia, WS; Natarajan, Pradeep, Broad Institute,

Cambridge, Massachusetts, 02142, US; Nekhai, Sergei, Howard University, Washington, District of

Columbia, 20059, US; Nelson, Sarah C., University of Washington, Biostatistics, Seattle, Washington,

98195, US; Neltner, Bonnie, University of Colorado at Denver, Denver, Colorado, 80204, US; Nessner,

Caitlin, Baylor College of Medicine Human Genome Sequencing Center, Houston, Texas, 77030, US;

Nickerson, Deborah, University of Washington, Department of Genome Sciences, Seattle, Washington,

98195, US; Nkechinyere, Osuji, Baylor College of Medicine Human Genome Sequencing Center,

Houston, Texas, 77030, US; North, Kari, University of North Carolina, Chapel Hill, North Carolina, 27599, US; O'Connell, Jeff, University of Maryland, Balitmore, Maryland, 21201, US; O'Connor, Tim,

University of Maryland, Baltimore, Maryland, 21201, US; Ochs-Balcom, Heather, University at Buffalo,

Buffalo, New York, 14260, US; Okwuonu, Geoffrey, Baylor College of Medicine Human Genome

Sequencing Center, Houston, Texas, 77030, US; Pack, Allan , University of Pennsylvania, Division of

Sleep Medicine/Department of Medicine, Philadelphia, Pennsylvania, 19104-3403, US; Paik, David T.,

Stanford University, Stanford Cardiovascular Institute, Stanford, California, 94305, US; Palmer,

Nicholette, Wake Forest Baptist Health, Biochemistry, Winston-Salem, North Carolina, 27157, US;

Pankow, James, University of Minnesota, Minneapolis, Minnesota, 55455, US; Papanicolaou, George,

National Heart, Lung, and Blood Institute, National Institutes of Health, Bethesda, Maryland, 20892, US;

Parker, Cora, RTI International, Biostatistics and Epidemiology Division, Research Triangle Park, North

Carolina, 27709-2194, US; Peloso, Gina, Boston University, Department of Biostatistics, Boston,

Massachusetts, 02118, US; Peralta, Juan Manuel, University of Texas Rio Grande Valley School of

Medicine, Edinburg, Texas, 78539, US; Perez, Marco, Stanford University, Stanford, California, 94305,

US; Perry, James, University of Maryland, Baltimore, Maryland, 21201, US; Peters, Ulrike, Fred

Hutchinson Cancer Research Center, Fred Hutch and UW, Seattle, Washington, 98109, US; Peyser,

Patricia, University of Michigan, Ann Arbor, Michigan, 48109, US; Phillips, Lawrence S, Emory

University, Atlanta, Georgia, 30322, US; Pleiness, Jacob, University of Michigan, Ann Arbor, Michigan,

48109, US; Pollin, Toni, University of Maryland, Baltimore, Maryland, 21201, US; Post, Wendy, Johns

Hopkins University, Cardiology/Medicine, Baltimore, Maryland, 21218, US; Powers Becker, Julia,

University of Colorado at Denver, Medicine, Denver, Colorado, 80204, US; Preethi Boorgula, Meher,

University of Colorado at Denver, Denver, Colorado, 80204, US; Preuss, Michael, Icahn School of

Medicine at Mount Sinai, New York, New York, 10029, US; Psaty, Bruce, University of Washington,

Seattle, Washington, 98195, US; Qasba, Pankaj, National Heart, Lung, and Blood Institute, National

Institutes of Health, Bethesda, Maryland, 20892, US; Qiao, Dandi, Brigham & Women's Hospital,

Boston, Massachusetts, 02115, US; Qin, Zhaohui, Emory University, Atlanta, Georgia, 30322, US;

Rafaels, Nicholas, University of Colorado at Denver, CCPM, Denver, Colorado, 80045, US; Raffield,

Laura, University of North Carolina, Genetics, Chapel Hill, North Carolina, 27599, US; Rajendran,

Mahitha, Baylor College of Medicine Human Genome Sequencing Center, Houston, Texas, 77030, US;

Ramachandran, Vasan S., Boston University, Boston, Massachusetts, 02215, US; Rao, D.C., Washington

University in St Louis, St Louis, Missouri, 63130, US; Rasmussen-Torvik, Laura, Northwestern

University, Chicago, Illinois, 60208, US; Ratan, Aakrosh, University of Virginia, Charlottesville,

Virginia, 22903, US; Redline, Susan, Brigham & Women's Hospital, Medicine, Boston, Massachusetts,

02115, US; Reed, Robert, University of Maryland, Baltimore, Maryland, 21201, US; Reeves, Catherine,

New York Genome Center, New York Genome Center, New York City, New York, 10013, US; Regan,

Elizabeth, National Jewish Health, Denver, Colorado, 80206, US; Reiner, Alex, Fred Hutchinson Cancer

Research Center, University of Washington, Seattle, Washington, 98109, US; Reupena, Muagututi‘a

Sefuiva, Lutia I Puava Ae Mapu I Fagalele, Apia, WS; Rice, Ken, University of Washington, Seattle,

Washington, 98195, US; Rich, Stephen, University of Virginia, Charlottesville, Virginia, 22903, US;

Robillard, Rebecca, University of Ottawa, Sleep Research Unit, University of Ottawa Institute for Mental

Health Research, Ottawa, ON K1Z 7K4, CA; Robine, Nicolas, New York Genome Center, New York

City, New York, 10013, US; Roden, Dan, Vanderbilt University, Medicine, Pharmacology, Biomedicla

Informatics, Nashville, Tennessee, 37235, US; Roselli, Carolina, Broad Institute, Cambridge,

Massachusetts, 02142, US; Rotter, Jerome, Lundquist Institute, Pediatrics, Torrance, California, 90502,

US; Ruczinski, Ingo, Johns Hopkins University, Baltimore, Maryland, 21218, US; Runnels, Alexi, New

York Genome Center, New York City, New York, 10013, US; Russell, Pamela, University of Colorado at

Denver, Denver, Colorado, 80204, US; Ruuska, Sarah, Blood Works Northwest, Seattle, Washington,

98104, US; Ryan, Kathleen, University of Maryland, Baltimore, Maryland, 21201, US; Sabino, Ester

Cerdeira, Universidade de Sao Paulo, Faculdade de Medicina, Sao Paulo, 01310000, BR; Saleheen,

Danish, Columbia University, New York, New York, 10027, US; Salimi, Shabnam, University of

Maryland, Pathology , Seattle, Washington, 98195, US; Salvi, Sejal, Baylor College of Medicine Human

Genome Sequencing Center, Houston, Texas, 77030, US; Salzberg, Steven, Johns Hopkins University, Baltimore, Maryland, 21218, US; Sandow, Kevin, Lundquist Institute, TGPS, Torrance, California,

90502, US; Sankaran, Vijay G., Harvard University, Division of Hematology/Oncology, Boston,

Massachusetts, 02115, US; Santibanez, Jireh, Baylor College of Medicine Human Genome Sequencing

Center, Houston, Texas, 77030, US; Schwander, Karen, Washington University in St Louis, St Louis,

Missouri, 63130, US; Schwartz, David, University of Colorado at Denver, Denver, Colorado, 80204, US;

Sciurba, Frank, University of Pittsburgh, Pittsburgh, Pennsylvania, 15260, US; Seidman, Christine ,

Harvard Medical School, Genetics, Boston, Massachusetts, 02115, US; Seidman, Jonathan, Harvard

Medical School, Boston, Massachusetts, 02115, US; Sériès, Frédéric, Université Laval, Quebec City,

G1V 0A6, CA; Sheehan, Vivien, Emory University, Pediatrics, Atlanta, Georgia, 30307, US; Sherman,

Stephanie L., Emory University, Human Genetics, Atlanta, Georgia, 30322, US; Shetty, Amol, University

of Maryland, Baltimore, Maryland, 21201, US; Shetty, Aniket, University of Colorado at Denver,

Denver, Colorado, 80204, US; Sheu, Wayne Hui-Heng, Taichung Veterans General Hospital Taiwan,

Taichung City, 407, TW; Shoemaker, M. Benjamin, Vanderbilt University, Medicine/Cardiology,

Nashville, Tennessee, 37235, US; Silver, Brian, UMass Memorial Medical Center, Worcester,

Massachusetts, 01655, US; Silverman, Edwin, Brigham & Women's Hospital, Boston, Massachusetts,

02115, US; Skomro, Robert, University of Saskatchewan, Saskatoon, SK S7N 5C9, CA; Smith, Albert

Vernon, University of Michigan, , ; Smith, Jennifer, University of Michigan, Ann Arbor, Michigan,

48109, US; Smith, Josh, University of Washington, Seattle, Washington, 98195, US; Smith, Nicholas,

University of Washington, Epidemiology, Seattle, Washington, 98195, US; Smith, Tanja, New York

Genome Center, New York, New York, 10013, US; Smoller, Sylvia, Albert Einstein College of Medicine,

New York, New York, 10461, US; Snively, Beverly, Wake Forest Baptist Health, Biostatistical Sciences,

Winston-Salem, North Carolina, 27157, US; Snyder, Michael, Stanford University, Stanford, California,

94305, US; Sofer, Tamar, Brigham & Women's Hospital, Boston, Massachusetts, 02115, US;

Sotoodehnia, Nona, University of Washington, Seattle, Washington, 98195, US; Stilp, Adrienne M.,

University of Washington, Seattle, Washington, 98195, US; Storm, Garrett , University of Colorado at

Denver, Genomic Cardiology, Aurora, Colorado, 80045, US; Streeten, Elizabeth, University of Maryland,

Baltimore, Maryland, 21201, US; Su, Jessica Lasky, Brigham & Women's Hospital, Channing

Department of Medicine, Boston, Massachusetts, 02115, US; Sung, Yun Ju, Washington University in St

Louis, St Louis, Missouri, 63130, US; Sylvia, Jody, Brigham & Women's Hospital, Boston,

Massachusetts, 02115, US; Szpiro, Adam, University of Washington, Seattle, Washington, 98195, US;

Taliun, Daniel, University of Michigan, Ann Arbor, Michigan, 48109, US; Tang, Hua, Stanford

University, Genetics, Stanford, California, 94305, US; Taub, Margaret, Johns Hopkins University,

Baltimore, Maryland, 21218, US; Taylor, Kent D., Lundquist Institute, Institute for Translational

Genomics and Populations Sciences, Torrance, California, 90502, US; Taylor, Matthew, University of

Colorado Anschutz Medical Campus, Aurora, Colorado, 80045, US; Taylor, Simeon, University of

Maryland, Baltimore, Maryland, 21201, US; Telen, Marilyn, Duke University, Durham, North Carolina,

27708, US; Thornton, Timothy A., University of Washington, Seattle, Washington, 98195, US;

Threlkeld, Machiko, University of Washington, University of Washington, Department of Genome

Sciences, Seattle, Washington, 98195, US; Tinker, Lesley, Fred Hutchinson Cancer Research Center,

Cancer Prevention Division of Public Health Sciences, Seattle, Washington, 98109, US; Tirschwell,

David, University of Washington, Seattle, Washington, 98195, US; Tishkoff, Sarah, University of

Pennsylvania, Genetics, Philadelphia, Pennsylvania, 19104, US; Tiwari, Hemant, University of Alabama,

Biostatistics, Birmingham, Alabama, 35487, US; Tong, Catherine, University of Washington, Department

of Biostatistics, Seattle, Washington, 98195, US; Tracy, Russell, University of Vermont, Pathology &

Laboratory Medicine, Burlington, Vermont, 05405, US; Tsai, Michael, University of Minnesota,

Minneapolis, Minnesota, 55455, US; Vaidya, Dhananjay, Johns Hopkins University, Baltimore,

Maryland, 21218, US; Van Den Berg, David, University of Southern California, USC Methylation

Characterization Center, University of Southern California, California, 90033, US; VandeHaar, Peter,

University of Michigan, Ann Arbor, Michigan, 48109, US; Vrieze, Scott, University of Minnesota,

Minneapolis, Minnesota, 55455, US; Walker, Tarik, University of Colorado at Denver, Denver, Colorado,

80204, US; Wallace, Robert, University of Iowa, Iowa City, Iowa, 52242, US; Walts, Avram, University of Colorado at Denver, Denver, Colorado, 80204, US; Wang, Fei Fei, University of Washington, Seattle,

Washington, 98195, US; Wang, Heming, Brigham & Women's Hospital, Mass General Brigham, Boston,

Massachusetts, 02115, US; Wang, Jiongming, University of Michigan, , US; Watson, Karol, University of

California, Los Angeles, Los Angeles, California, 90095, US; Watt, Jennifer, Baylor College of Medicine

Human Genome Sequencing Center, Houston, Texas, 77030, US; Weeks, Daniel E., University of

Pittsburgh, Pittsburgh, Pennsylvania, 15260, US; Weinstock, Joshua, University of Michigan,

Biostatistics, Ann Arbor, Michigan, 48109, US; Weir, Bruce, University of Washington, Seattle,

Washington, 98195, US; Weiss, Scott T, Brigham & Women's Hospital, Channing Division of Network

Medicine, Department of Medicine, Boston, Massachusetts, 02115, US; Weng, Lu-Chen, Massachusetts

General Hospital, Boston, Massachusetts, 02114, US; Wessel, Jennifer, Indiana University,

Epidemiology, Indianapolis, Indiana, 46202, US; Willer, Cristen, University of Michigan, Internal

Medicine, Ann Arbor, Michigan, 48109, US; Williams, Kayleen, University of Washington, Biostatistics,

Seattle, Washington, 98195, US; Williams, L. Keoki, Henry Ford Health System, Detroit, Michigan,

48202, US; Wilson, Carla, Brigham & Women's Hospital, Boston, Massachusetts, 02115, US; Wilson,

James, Beth Israel Deaconess Medical Center, Cardiology, Cambridge, Massachusetts, 02139, US;

Winterkorn, Lara, New York Genome Center, New York City, New York, 10013, US; Wong, Quenna,

University of Washington, Seattle, Washington, 98195, US; Wu, Joseph, Stanford University, Stanford

Cardiovascular Institute, Stanford, California, 94305, US; Xu, Huichun, University of Maryland,

Baltimore, Maryland, 21201, US; Yanek, Lisa, Johns Hopkins University, Baltimore, Maryland, 21218,

US; Yang, Ivana, University of Colorado at Denver, Denver, Colorado, 80204, US; Yu, Ketian,

University of Michigan, Ann Arbor, Michigan, 48109, US; Zekavat, Seyedeh Maryam, Broad Institute,

Cambridge, Massachusetts, 02142, US; Zhang, Yingze, University of Pittsburgh, Medicine, Pittsburgh,

Pennsylvania, 15260, US; Zhao, Snow Xueyan, National Jewish Health, Denver, Colorado, 80206, US;

Zhao, Wei, University of Michigan, Department of Epidemiology, Ann Arbor, Michigan, 48109, US;

Zhu, Xiaofeng, Case Western Reserve University, Department of Population and Quantitative Health

Sciences , Cleveland, Ohio, 44106, US; Zody, Michael, New York Genome Center, New York, New

York, 10013, US; Zoellner, Sebastian, University of Michigan, Ann Arbor, Michigan, 48109, US
